# Nonribosomal peptide synthetases require dynamic interaction between modular domains

**DOI:** 10.1101/2023.11.13.566838

**Authors:** Ye-Jun Peng, Xiaoli Zeng, Yuxing Chen, Cong-Zhao Zhou, Wei Miao, Yong-Liang Jiang, Cheng-Cai Zhang

## Abstract

Nonribosomal peptide synthetases (NRPSs) are large multidomain enzymes for the synthesis of a variety of bioactive peptides in a modular and pipelined fashion. Here, we investigated how the condensation (C) domain and the adenylation (A) domain cooperate with each other for the efficient catalytic activity in microcystin NRPS modules. We solved two crystal structures of the microcystin NRPS modules, representing two newly identified conformations in the NRPS catalytic cycle. Our data reveals that the dynamic interaction between the C and the A domains in these modules are mediated by the conserved “RXGR” motif, and this interaction is important for the adenylation activity. Furthermore, the “RXGR” motif-mediated dynamic interaction and its functional regulation is prevalent in different NRPSs modules possessing both the A and the C domains. This study provides new insight into the catalytic mechanism of NRPSs and should inspire novel ideas in NRPS enzyme engineering in synthetic biology.

## Introduction

Nonribosomal peptide synthetases (NRPSs) are mega-enzymes that can assemble proteinogenic and nonproteinogenic amino acids into a vast variety of peptides, named nonribosomal peptides (NRPs) (Strieker *et al*, 2010). Many NRPs are therapeutic agents, with antibacterial, antiviral, immunosuppressor, or antitumor activities (Felnagle *et al*, 2008). NRPSs synthesize peptides in a modular and pipelined fashion, and each module adds one amino acid to the nascent peptide (Fischbach & Walsh, 2006; Koglin & Walsh, 2009). Therefore, the number and the specificity of the modules determine the length and the sequence of the amino acid in the corresponding peptide product.

A basic NRPS module for peptide elongation (elongation module) consists of a condensation (C) domain, an adenylation (A) domain, and a peptidyl carrier protein (PCP) domain (Strieker *et al*., 2010; Weber & Marahiel, 2001). The A domain is responsible for the selection and activation of the cognate amino acid, and then transfers the activated substrate to the preactivated phosphopantetheinyl (PPant) arm attached to the adjacent PCP domain (Crosby & Crump, 2012; Gulick, 2009). In the next step, the PCP domain forwards the covalently bound substrate to the C domain of the same module (Crosby & Crump, 2012). The C domain catalyzes the peptide bond formation between the substrate from the PCP domain of the current module and the peptide chain carried on the preceding module at the active site, leading to the elongation of the peptide chain (Wang *et al*, 2022). After condensation, the current PCP domain brings the elongated peptide to the C domain of the next module for another round of condensation reaction (Koglin & Walsh, 2009; Strieker *et al*., 2010). Usually, the initiation NRPS module in the assembly line lacks the C domain, and the termination module contains an extra thioesterase (TE) domain, which releases the peptide by cyclization or hydrolysis at the end of the whole synthesis (Lawson *et al*, 1994). Moreover, the diversity of NRPs may further be expanded by the action of specialized tailoring domains involved in the modification of the growing peptides, and such a domain may be part of a NRPS module or stand alone as a protein by itself (Konz & Marahiel, 1999; Süssmuth & Mainz, 2017; Walsh *et al*, 2001).

Biochemical and structural studies on the NRPS stand-alone domains (A, C, PCP) and NRPS didomains (A-PCP, PCP-C) indicate that the overall 3D structures of the A and C domains are conserved, but they display different conformations, suggesting that their structure changes dynamically during catalysis (Bloudoff *et al*, 2013; Conti *et al*, 1997; Mitchell *et al*, 2012; Patel *et al*, 2023; Reger *et al*, 2008; Samel *et al*, 2007; Tan *et al*, 2015). The A domain consists of a large A_core_ (∼450 amino acid residues) subdomain at the N-terminus and a small A_sub_ (∼100 amino acid residues) subdomain at the C-terminus (Gulick, 2009; Mitchell *et al*., 2012; Tan *et al*., 2015). Ten conserved motifs, namely A1 to A10, were found to play important structural and catalytic roles in enzymatic reactions (Turgay *et al*, 1992). Three major conformations of the A domain have been reported (Yonus *et al*, 2008): (a) **an open conformation**, in which A_sub_ is rotated 30° away from the active site of A_core,_ in preparation for entry of ATP and Mg^2+^ (Tanovic *et al*, 2008); (b) **an adenylation (or closed) conformation**, in which the substrate and the cofactors are bound at the active site covered by A_sub_, allowing the adenylation reaction to occur (Reimer *et al*, 2016). (c) **a thiolation conformation**, in which A_sub_ rotates by ∼140° around A_core_, to facilitate the attachment of the adenylated substrate to the PPant arm of the PCP domain by thiolation (Reger *et al*., 2008; Yonus *et al*., 2008). The C domain (∼450 amino acids) adopts two chloramphenicol acetyltransferase folds, with the N- and C-terminal subdomains (C_N_ and C_C_) forming a “V-shaped” pseudo dimer. The key catalytic site (HHXXXDG motif) for condensation is located at the interface of the two subdomains. For the condensation reaction, the two amino acids charged on the donor and the acceptor domains of PCP are brought together to the key catalytic site through two catalytic tunnels (donor tunnel and acceptor tunnel), located in the front and the back sides of the “V-shaped” structure, respectively (Bloudoff *et al*., 2013; Keating *et al*, 2002). The C domain has been proposed to exist in two major conformations: (a) **an open conformation**, in which no substrate is bound at the two tunnels and the “V shaped” pseudo dimer is relatively open; (b) **a closed state or condensation conformation**, in which at least one of the substrates is inserted at one tunnel, and the V shaped pseudo dimer is relatively more closed than that found in the open state (Bloudoff *et al*., 2013; Keating *et al*., 2002).

The increasing number of crystal structures of multidomains, modules, and cross-modules of NRPSs reported in different catalytic states highlights the importance of the dynamic interactions of the different domains during the catalytic cycle of NRPSs (Izore & Cryle, 2018; Patel *et al*., 2023; Wang *et al*., 2022). The PCP domain was found to interact with different catalytic domains at different regions during catalysis, and mutation of conserved residues in the observed interaction interfaces have all resulted in a loss or a reduction of the catalytic activity (Drake *et al*, 2006; Frueh *et al*, 2008; Koglin *et al*, 2006; Samel *et al*., 2007). In addition to the dynamic domain-domain interactions related to PCP domains during catalysis, flexible interaction between adjacent A and C domains has also been found in different NRPSs. In the module or di-module crystal structures of SrfA-C (Tanovic *et al*., 2008), AB3403 (Drake *et al*, 2016), ObiF (Kreitler *et al*, 2019) and LgrA (Reimer *et al*, 2019), each C domain shares an extensive interaction surface with the adjacent and cognate A domain. In contrast, in the EntF termination module, the interaction surface between the A and the C domains is much reduced as compared to that found in others (Drake *et al*., 2016; Miller *et al*, 2016a). The reason behind such a difference in the interaction interfaces lie in the fact that these structures are in different catalytic states. The rotation of the PCP domain and the A_sub_ domain is necessary for delivery of the peptide intermediates to the different catalytic domains (Reimer *et al*., 2016; Tanovic *et al*., 2008). Recently, a few studies on NRPS modules involved in microcystin or sulfazecin production indicate that the co-existence of the adjacent A and C domains in some NRPS modules is necessary for substrate selectivity or the adenylation activity of the A domain (Kaniusaite *et al*, 2019; Li *et al*, 2017; Meyer *et al*, 2016). These suggest the importance of the C domain in controlling dynamic domain-domain interactions in the conformational space of the flexible A_sub_-and PCP domains (Meyer *et al*., 2016). However, the underlying molecular mechanism remains poorly understood.

Microcystins (MCs) are typical NRPs produced by a diverse range of bloom-forming freshwater cyanobacteria, including those belonging to the genera of *Microcystis*, *Planktothrix*, *Nostoc* and *Oscillatoria* (Duy *et al*, 2000; Rinehart *et al*, 1994; van Apeldoorn *et al*, 2007). MCs are the most common hepatotoxins present in aquatic environment, which constitute a great threat to human and animal health (Massey *et al*, 2022). To date, more than 270 MC variants with varying levels of toxicity have been identified (Massey *et al*., 2022), and all of them share a common cyclic heptapeptide structure (D-Ala^1^-X^2^-D-MeAsp^3^/D-Asp^3^-Z^4^-Adda^5^-D-Glu^6^-Mdha^7^), in which X and Z represent variable amino acids at position 2 and 4, D-MeAsp^3^ is D-erythro-β-methyl aspartic acid, Mdha^7^ is N-methyl dehydroalanine, and Adda^5^ is 3-amino-9-methoxy-2,6,8-trimethyl-10-phenyldeca-4,6-dienoic acid, respectively (Botes *et al*, 1984; Rinehart *et al*., 1994). In *Microcystis aeruginosa* PCC 7806 (here-after *Microcystis*), MCs are predicted to be synthesized by a megaenzyme complex encoded by a 55-kb long gene cluster, the *mcy* gene cluster (Figure 1A) (Dittmann *et al*, 1997; Meiβner *et al*, 1996; Tillett *et al*, 2000). The corresponding proteins McyA, McyB and McyC are NRPSs (Figure 1B), while McyD is a PKS (polyketide synthase), and McyE and McyG are PKS/NRPS hybrids. Four proteins, McyF (amino acid racemase), McyH (ABC transporter), McyI (putative dehydrogenase), and McyJ (O-methyltransferase) are tailoring enzymes (Tillett *et al*., 2000). Based on the domain organization of these proteins in comparison with those of other PKSs and NPRSs, a biosynthetic pathway of MCs has been proposed (Tillett *et al*., 2000). *Microcystis* produces predominantly MC-LR and MC-(D-Asp^3^)-LR, containing leucine (L) at position 2 and arginine (R) at position 4 (Figure 1C) (Meyer *et al*., 2016). The synthesis of MCs starts at the Adda^5^ residue. Both McyA (McyA_1_ and McyA_2_ modules) and McyB (McyB_1_ and McyB_2_ modules) consist of two NRPS modules, and are predicted to add, sequentially, amino acids Mdha^7^, D-Ala^1^, L-Leu^2^, D-MeAsp^3^/D-Asp^3^ during MC synthesis (Figure 1B) (Tillett *et al*., 2000). Since the A domain of the McyA_2_ module is predicted to activate L-alanine (L-Ala), the epimerization domain (tailoring domain) at the C-terminus of this module was likely to be responsible for the modification of the bound L-Ala to D-Ala (Tillett *et al*., 2000). McyC is the terminal NRPS module responsible for adding the amino acid L-Arg onto the elongated linear peptide chain, and cyclizing and releasing the peptide by the thioesterase domain (TE) at the C-terminal end (Tillett *et al*., 2000).

**Figure 1.**
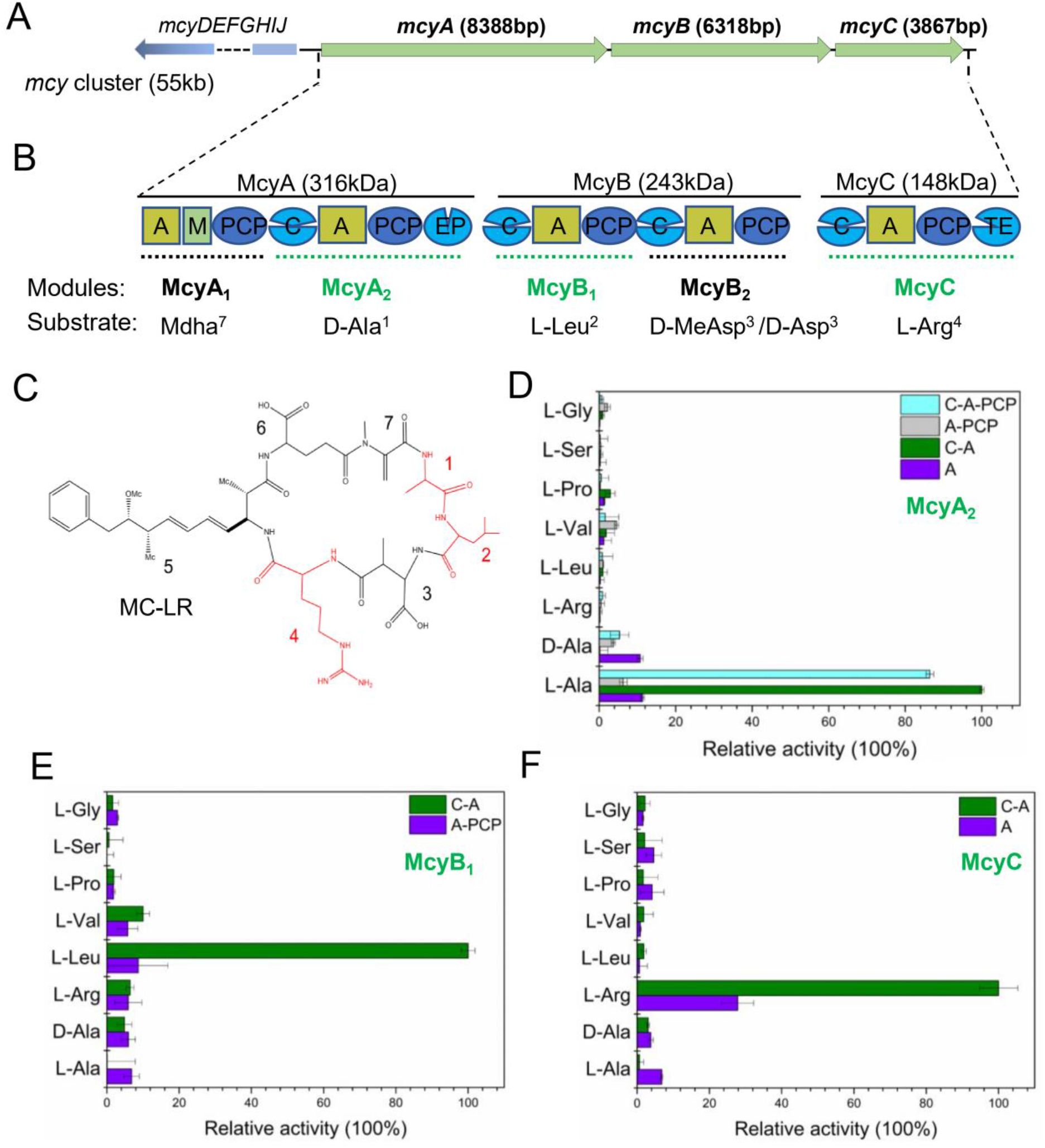
The presence of the C domain enhances the adenylation activity of the A domain in three MC NRPS modules. (A) Schematic representation of the MC biosynthetic gene cluster (*mcy*) in *Microcystis*. (B) Domain organization of McyA, McyB, McyC. A: adenylation domain, M: N-Methyltransferase domain, PCP: peptidyl carrier protein domain, C: condensation domain, EP: Epimerization domain, TE: Thioesterase domain. The NRPS modules are marker by dashed lines, and those characterized in this study by green dashed lines. The name of each module and the corresponding substrate predicted for these modules are indicated below the domain structure. (C) Chemical structure of the cyclo-heptapeptide microcystin-LR, and the sites 1, 2 and 4 are highlighted in red. (D) In vitro adenylation activities of different truncated polypeptides of McyA_2_. (E) In vitro adenylation activities of different truncated polypeptides of McyB_1_. (F) In vitro adenylation activities of different truncated polypeptides of McyC. Adenylation activities are determined using a continuous hydroxylamine release assay (see Materials and Methods for details). The OD_650_ value of the WT protein is set to 100%. Error bars indicate SD from the mean of two experimental replicates.

In this study, we found that the presence of the C domain significantly promoted the activity of the cognate A domain in vitro for three MC NRPS modules. Crystal structural analysis, mutagenesis of key residues and cysteine-based crosslinking studies demonstrated that the dynamic interaction between the C and the A domains during the catalytic cycle of McyA_2_ and McyB_1_ modules are mediated by the “RXGR” motif and is important for the adenylation activity of the cognate A domain. Such a mechanism is prevalent in elongation and termination modules belonging to different NRPSs. These findings provide insight into the catalytic mechanism of NRPSs.

## Results

### The C domain enhances the adenylation activity of the cognate A domain in three MC NRPS modules

A previous study suggested that the A domains of McyB_1_ and McyC modules show the anticipated substrate activation in vitro only in the presence of its cognate C domain (Meyer *et al*., 2016). To further investigate the relationship between the C and the A domain in other MC NRPS modules, different modules, McyA_1_, McyA_2_, McyB_1_, McyB_2_ and McyC, were expressed in *Escherichia coli* in different combination with single A domains, C-A and A-PCP bidomains, and C-A-PCP tri-domains, respectively (Figure S1A). The recombinant proteins were purified by Ni-NTA affinity chromatography followed by size exclusive chromatography. At the end, the single A domains from McyA_2_ and McyC modules, the C-A bidomains from McyA_2_, McyB_1_ and McyC modules, the A-PCP bidomains from McyA_2_ and McyB_1_ modules, and the C-A-PCP tri-domain of McyA_2_ module could be expressed and purified (Figure S1B). The adenylation activity of the purified proteins was tested by a continuous hydroxylamine release assay (Figure S2), using different amino acids as substrates. As shown in Figure 1 (panels D-F), the C-A bidomains of McyA_2_, McyB_1_ and McyC modules could activate the corresponding specific substrate with the strongest activity (L-Ala for McyA_2_, L-Leu for McyB_1_, L-Arg for McyC), consistent with the previous prediction (Meyer *et al*., 2016; Tillett *et al*., 2000). However, the single A domains or A-PCP bidomains from the same three modules displayed a much weaker adenylation activity than the C-A bidomain proteins towards the same substrates (Figure 1D-1F). The results related to the McyB_1_ and McyC modules were thus consistent with those reported (Meyer *et al*., 2016). These results indicated that in addition to McyB_1_ and McyC modules, the activity of the A domain in McyA_2_ modules also depends on the presence of the cognate C domain. Furthermore, we analyzed the adenylation activity of both the A-PCP bidomain and the C-A-PCP tri-domain polypeptides from the McyA_2_ module. We found that McyA_2_-(C-A-PCP) also showed much stronger adenylation activity than McyA_2_-(A-PCP) (Figure 1D). Taken together, we concluded that the presence of the C domain enhances the activity of the cognate A domain in MC NRPS modules.

### Crystal structures of McyA_2_-(C-A-PCP) and McyB_1_-(C-A)

To gain deeper insight into the relationship between the C and A domains during catalysis in MC NRPS modules, we performed crystallization assays, and finally solved the crystal structure of McyA_2_-(C-A-PCP) and McyB_1_-(C-A) in the presence of the corresponding substrate L-Ala/Mg·ATP and L-Leu/Mg·ATP, with 2.45-Å and 2.70-Å resolutions, respectively.

The overall structure of McyB_1_-(C-A) consists of an N-terminal C domain (residues S14-T450), and a C-terminal A domain (S473-D963) (Figure 2A). Similarly, McyA_2_-(C-A-PCP) can be divided into an N-terminal C domain (residues Q1269-L1709), an A domain (residues C1732-N2224), and a C-terminal PCP domain (residues E2237-T2306) that is positioned near the acceptor tunnel of the C domain (Figure 2B). The overall folds of the individual C, A and PCP domains from these two structures were consistent with previously reported structures (Izore & Cryle, 2018; Koglin & Walsh, 2009; Patel *et al*., 2023). In both structures, the C domain adopts two chloramphenicol acetyltransferase folds, with the N- and the C-terminal subdomains forming a typical “V-shaped” pseudo dimer. The active site harboring the conserved HHXXXD motif is located on a loop between the β7 strand and the α4 helix of each C domain (Figure 2A and 2B). Each A domain is subdivided into an A_core_ and an A_sub_ subdomains. Both A_core_ subdomains are composed of two αβαβα regions that form a sandwich and a β-barrel, respectively (Figure 2A and 2B). The A_sub_ subdomain in McyB_1_-(C-A) is comprised of a central three-stranded β-sheet (β37-β39) surrounded by two helices (α26-α27) (Figure 2A), whereas only two β-sheets (β37-β38) and one helix (α27) can be well modeled in the McyA_2_-(C-A-PCP) structure due to the weak electron density in this region (Figure 2B). The PCP domain in McyA_2_-(C-A-PCP) consists of a four helical bundle (α29-α32), and the conserved residue S2267 for attachment of the phosphopantetheinyl (PPant) arm is located at the start of α29 (Figure 2B).

**Figure 2.**
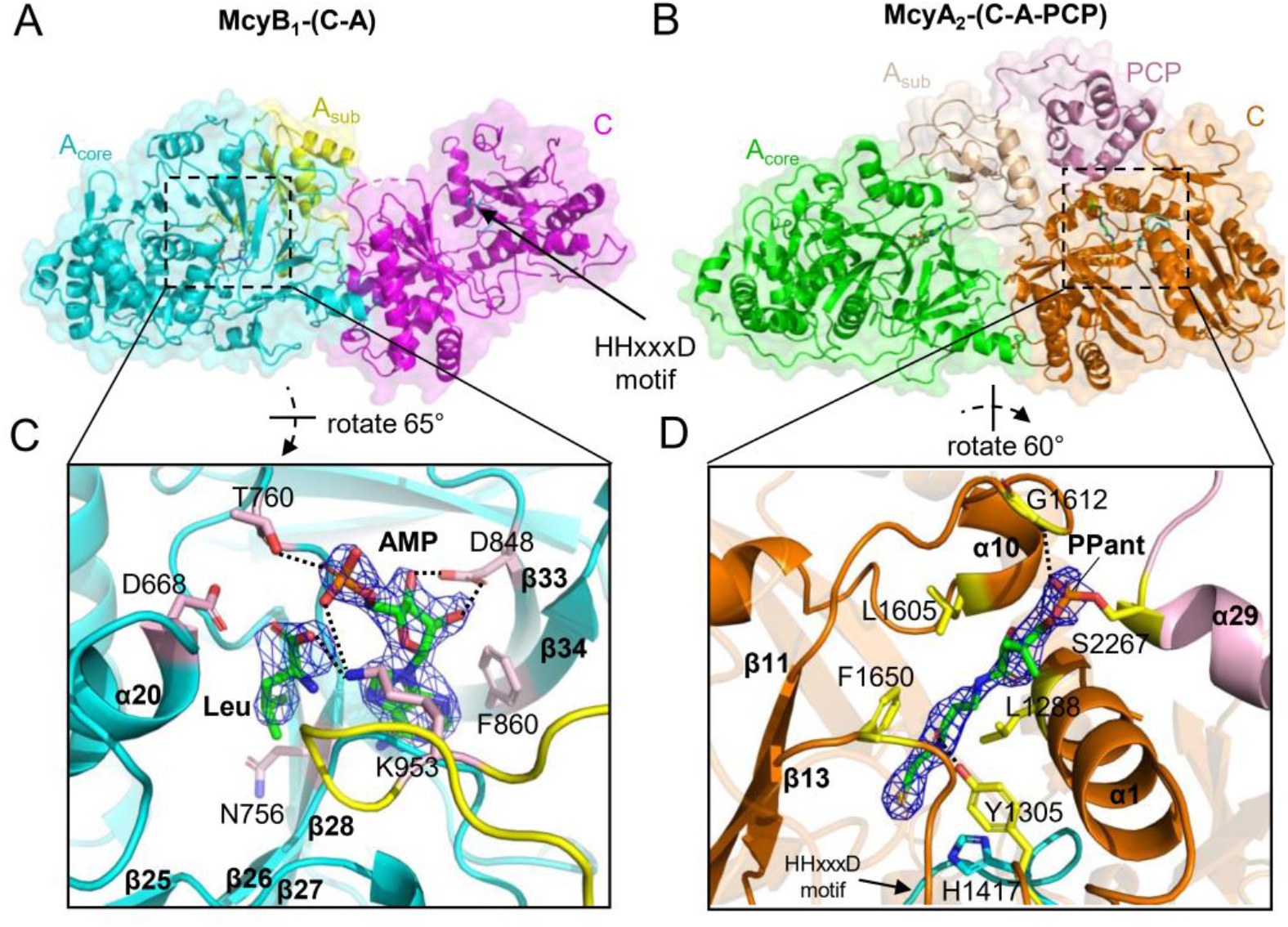
Overall crystal structures of McyB_1_-(C-A) and McyA_2_-(C-A-PCP). (A) Ribbon structure of McyB_1_-(C-A), with the C domain in magenta, the A_core_ domain in cyan, and the A_sub_ domain in yellow. One Leu and one molecule of AMP located at the active site of the A domain are shown in green/orange-colored sticks, respectively. The “HHXXXD” motif (active site) of the C domain is indicated by black arrow. (B) Ribbon structure of McyA_2_-(C-A-PCP), with C domain in orange, A_core_ domain in green, A_sub_ domain in wheat, and PCP domain in light pink. One molecule of ATP in the A domain, a PPant arm inserted in the C domain are shown in green/orange-colored sticks. The “HXXXD” motif in the C domain is indicated by cyan. (C) Details of binding pockets for Leu and AMP in McyB_1_-(C-A). The Leu residue and AMP are shown in green/orange-colored sticks, superimposed with the *Fo-Fc* map contoured at 2σ. The key polar interactions to the residues are indicated by dashed lines. The residues interacting with Leu and AMP are shown in stick representation with pink color. (D) Interaction of PPant arm in C domain of McyA_2_-(C-A-PCP). The PPant arm is shown in green/orange-colored sticks and are superimposed with the *Fo-Fc* map contoured at 2σ. The PPant arm is covalently attached to Ser2267 of the PCP domain (light pink). The residues interacting with the PPant arm are shown in stick representation with yellow color and the key polar interactions are indicated by dashed line.

In the structure of McyB_1_-(C-A), the electron density corresponding to L-Leu and AMP was clear in the pocket of the A domain. The sites for specific binding of L-Leu in the A domain include D668 from α20, N756 from β27, and K953 from A10 catalytic loop. (Figure 2C and S3). The AMP is held in place by residues T760 from η10-α26 loop, D848 from β33, F860 from β34 and K953. The conserved K953 interacts with the oxygen atom from the phosphate group in AMP and the carboxylate group in L-Leu, respectively (Figure 2C). In McyA_2_-(C-A-PCP), some weak electron density could be found near the active site of the A domain, and an ATP molecule fits well here (Figure S4A). Moreover, a clear electron density was present corresponding to the PPant arm inserted into the acceptor tunnel of the C domain of McyA_2_-(C-A-PCP) on the one hand, and covalently attached to the conserved S2267 residue from the PCP domain on the other (Figure 2D). The wall of the acceptor tunnel is formed by residues L1288 from α1, Tyr1305 from the linker between α1 and β2, F1650 from the loop after β13, and L1605 from α10. The interactions between the pantetheine cofactor and the side chain of these residues hold the PPant arm close to the active site (HHXXXD motif) (Figure 2D).

### The structures of McyA_2_-(C-A-PCP) and McyB_1_-(C-A) represent two distinct NRPS conformations

The structural variations of McyA_2_-(C-A-PCP) and McyB_1_-(C-A) suggest that they present different conformations in the NRPS catalytic cycle. To confirm this observation, we further compared these structures with two previously well characterized NRPSs, SrfA-C (Tanovic *et al*., 2008) and AB3403 (Drake *et al*., 2016). Comparison of the four structures reveled that the C domain in McyA_2_-(C-A-PCP) adopts a “closed” conformation similar as that of AB3403, in which the N- and C-terminal subdomains are close to each other and the PPant arm is present in the active site (Figure 3A). In contrast, the C domain in McyB_1_-(C-A) adopts an “open” conformation similar as that of SrfA-C, in which no substrate binding occurred at the tunnels, and the V shaped pseudo dimers are in an open state (Figure 3A).

**Figure 3.**
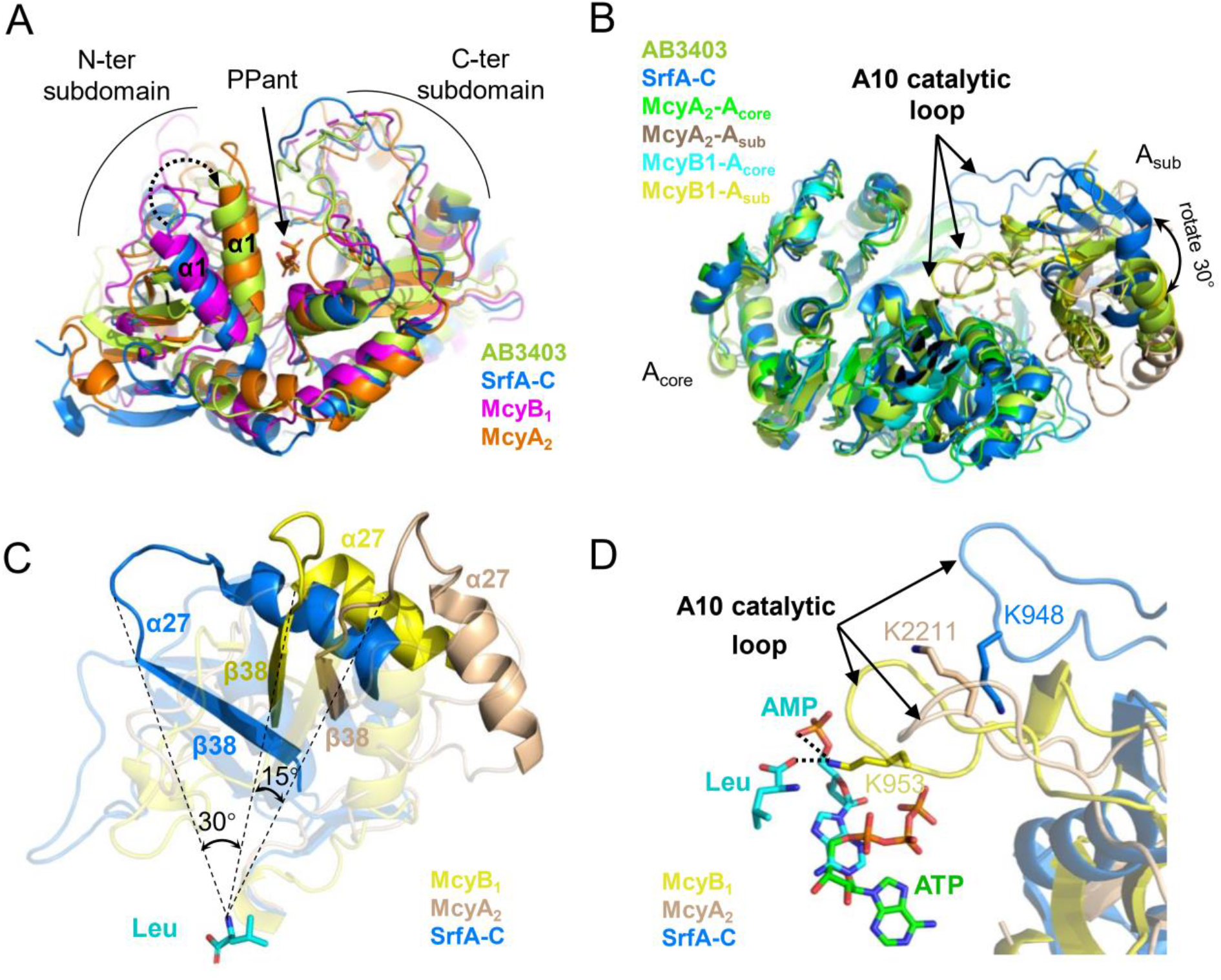
Structural analysis of McyB_1_-(C-A) and McyA_2_-(C-A-PCP). (A) Superposition of the C domains of McyB_1_ (magenta), McyA_2_ (orange), SrfA-C (marine, PDB: 2VSQ) and AB3403 (limon, PDB: 4ZXH). (B) Superposition of the A domains of McyB_1_ (A_core_, in cyan; A_sub_, in yellow), McyA_2_ (A_core_, in green; A_sub_, in wheat), SrfA-C (in marine) and AB3403 (in limon). (C) Rotation angle of the A_sub_ domains of McyB_1_ (in yellow), McyA_2_ (in wheat) and SrfA-C (in marine), relative to their respective active site (Leu as coordinate). The β38 and α27 regions of the A_sub_ domains of the three structures are highlighted. (D) Close-up view of the A10 catalytic loop of McyB_1_ (in yellow), McyA_2_ (in wheat) and SrfA-C (in marine) in the structural alignment. The conserved Lys residue in the A10 catalytic loop, the ATP molecule (green-orange) in McyA_2_, and the Leu residue and the AMP molecule (cyan-orange) in McyB_1_ are shown in stick representation. The interactions between the K953 residue and the ligands (Leu and AMP) in McyB_1_ are indicated by dashed lines.

When the A domains of these structures were aligned, the A_core_ subdomains could be well superimposed with each other, but the difference in the relative positions of the A_sub_ subdomains are noticeable (Figure 3B). The A_sub_ in McyB_1_-(C-A) is closed upon the active site and well aligned with that of AB3403 (Figure 3B), indicating the A domain of McyB_1_-(C-A) adopts the same “adenylation” conformation (or “closed” conformation) as AB3403. In contrast, the A domain of SrfA-C is in an “open” conformation, whereby its A_sub_ is oriented 30° away from the active site (Tanovic *et al*., 2008). In the case of McyA_2_-(C-A-PCP), its A domain displays an overall “closed” conformation similar to those found in the structures of McyB_1_-(C-A) and AB3403. However, a close-up view of this structure indicates an important rotational offset as compared to other “closed” conformation structures (Figure 3B and S4B). These results suggest that the A domain in the structure of McyA_2_-(C-A-PCP) could correspond to an intermediate conformation.

Indeed, by comparing the relative positions of all the A_sub_ in the aligned structures, we observed a rotation by 45° relative to the catalytic sites between SrfA-C and McyA_2_-(C-A-PCP), instead of a rotation by 30° observed between the SrfA-C (“open” conformations) and McyB_1_-(C-A) (“adenylation” conformation). In other words, the A_sub_ of McyA_2_-(C-A-PCP) is rotated by further 15° in comparison to that of McyB_1_-(C-A) (Figure 3C). As a consequence, the position of the critical A10 catalytic loop in the A_sub_ of McyA_2_-(C-A-PCP) is also different from those of SrfA-C and McyB_1_-(C-A) (Figure 3D). In SrfA-C, the conserved K948 in the A10 catalytic loop moves away from the active site so that ATP and the substrate can be loaded to the active site (Tanovic *et al*., 2008). In McyB_1_-(C-A), the conserved K953 positions in close proximity to the carboxylate group of the L-Leu and the phosphate group of AMP, providing a catalytic impetus for the adenylation reaction. However, in McyA_2_-(C-A-PCP), the conserved residue K2211 is located in a position far away from the active site, and no substrates are present at the binding pocket, despite the fact that the A10 catalytic loop is close to the active site. Moreover, the ATP ligand is observed in a reversed conformation near, but not within the ATP binding pocket, which is occupied by AMP in the structure of McyB_1_-(C-A) (Figure 3D and Figure S4A). These observations suggest that the structure of the A domain in McyA_2_-(C-A-PCP) corresponds to an intermediate conformation between the reported “open” and “closed” conformations, in a transition from the former to the latter. Therefore, we refer to this intermediate conformation as the “pre-adenylation” conformation, which corresponds to the inception of ATP binding prior to the catalytic reaction. Compared to all published NRPS structures containing both the A and C domains (Drake *et al*., 2016; Kreitler *et al*., 2019; Reimer *et al*., 2019; Tanovic *et al*., 2008), the conformational combination of the A and C domains in the structures of McyA_2_-(C-A-PCP) and McyB_1_-(C-A) represent thus two different NRPS conformations (Figure S4C).

### Dynamic interaction between the C and A domains enhances the adenylation activity of McyA_2_-(C-A-PCP) and McyB_1_-(C-A)

In the structures of McyA_2_-(C-A-PCP) and that of McyB_1_-(C-A), two different interfaces between the C and the A domains are observed, referred to as the top and the bottom interfaces, respectively (Figure 4A). For the bottom interface, the β30-β31 loop of the A_core_ subdomain is stacked against the α11 and the α8-α9 loop of the C domain in both structures. Similar to other NRPSs (Reimer *et al*., 2016; Tanovic *et al*., 2008), this bottom interface forms a stable platform on which the PCP domain and the A_sub_ domain can rearrange to promote the catalytic cycle. Although the numerous residues involved in the interaction are poorly conserved, they form superimposable structures (Figure 4A and Figure S3). In the top interfaces, the interactions involve the η10-α26 loop of the A_sub_ domain, and the α10-β11 loop together with a stretch of residues after α6 in the C domain. Although the sequences of the η10-α26 loop shows a high level (67%) of similarity, the interfaces in these two structures are spatially different due to the rotation of the η10-α26 loop (Figure 4A, Figure S3 and S5). In McyA_2_-(C-A-PCP), two conserved residues, R2129 and R2132 of the “RGXR” motif from the η10-α26 loop, play important roles in stabilizing the top interface by forming hydrogen bonds with Q1613 and V1482, and one salt bridge with D1481, respectively (Figure 4B and Figure S3). In contrast, in McyB_1_-(C-A), only one salt bridge is formed between the conserved R874 and E359 at the top interface (Figure 4C).

**Figure 4.**
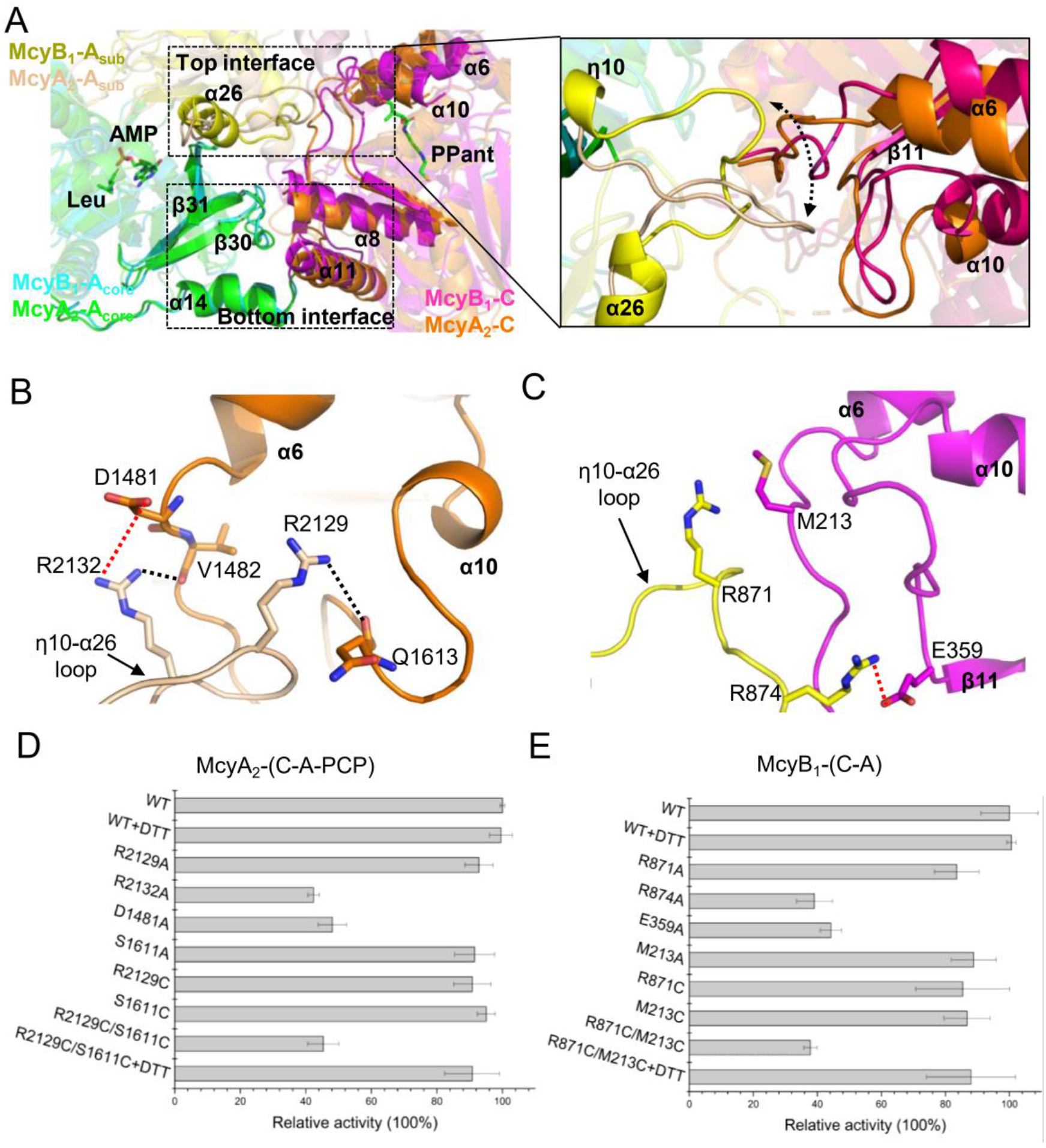
Structural and functional characterization of the interaction between the A and the C domains in McyB_1_-(C-A) and McyA_2_-(C-A-PCP). (A) Comparison of the interaction interface between the C domain (McyB_1_-C, in magenta; McyA_2_-C, in orange) and the A domain (McyB_1_-A_core_, in cyan; McyB_1_-A_sub_, in yellow; McyA_2_-A_core_, in green; McyA_2_-A_sub_, in wheat) in McyB_1_-(C-A) and McyA_2_-(C-A-PCP). In the left panel, two interaction interfaces between the C domain and the A domain in these structures are boxed and labeled as “bottom interface” and “top interface”, respectively. In the right panel, a zoom-in view of the difference at the “top interface” in McyB_1_-(C-A) and McyA_2_-(C-A-PCP). The rotation of the η10-α26 loop of the A_sub_ domain in McyB_1_-(C-A) or McyA_2_-(C-A-PCP) is indicated by a dotted line with double arrows. (B-C) Direct contact interactions between the η10-α26 loop and the C domain in the McyA_2_-(C-A-PCP) structure (B) and the McyB_1_-(C-A) structure (C). The salt bridge is indicated by a red dashed line, and the hydrogen bond is indicated by a black dashed line. The key residues involved in the interactions are shown in stick representation. (D-E) The adenylation activation profile of McyA_2_-(C-A-PCP) (D) and McyB_1_-(C-A) (E) toward the respective specific substrate (L-Leu for McyB_1_-(C-A), L-Ala for McyA_2_-(C-A-PCP)) with different treatments, including single point mutation, crosslink by double Cys substitution. Adenylation activities are determined using a continuous hydroxylamine release assay. The OD_650_ value of the WT protein is set at 100%. Error bars indicate standard deviation based on the mean of two experimental replicates.

To evaluate the functional importance of the obseved interactions between the C and A_sub_ domains in McyA_2_-(C-A-PCP) and McyB_1_-(C-A), we substituted each of the two conserved Arg residues by Ala in the “RGXR” motif and tested the effect of these mutations on the adenylation activity. As shown in Figure 4D and 4E, these mutants all exhibited reductions in their adenylation activity, especially for McyA_2_-(C-A^R2132A^-PCP) and McyB_1_-(C-A^R874A^), both of which had a reduction by 60% as compare to the wild-type proteins. To rule out the possibility that such an effect was caused by a functional disruption of the A domain in McyA_2_-(C-A^R2132A^-PCP) and McyB_1_-(C-A^R874A^) mutants, we generated the variants McyA_2_-(C^D1481A^-A-PCP) and McyB_1_-(C^E359A^-A), which disrupted the same interaction, but by changing the residues on the C domains. A similar reduction of the activity using specific substrate was also observed (Figure 4D and 4E). These results indicated that the optimal adenylation activity of the A domain in McyA_2_-(C-A-PCP) and McyB_1_-(C-A) depends on the interactions at the top interface between the C and A_sub_ domains.

We further examined the function of the interaction between the C and A domains of McyA_2_-(C-A-PCP) and McyB_1_-(C-A) by fixing their interface using a Cys-based cross-linking approach. To achieve an efficient interface crosslinking, a pair of residues located at the top interface in close proximity are selected and mutated into Cys. The R2129C/S1611C pair in McyA_2_-(C-A-PCP) and the R871C/M213C pair in McyB_1_-(C-A) were selected for the experiments (Figure 4D and 4E). When these residues were individually substituted by either alanine or cysteine, the corresponding protein variant exhibited only a marginal reduction in their adenylation activity (8-12% reduction as compared to the WT). However, when the pair of residues were both replaced by Cys, the corresponding proteins McyA_2_-(C^S1611C^-A^R2129C^-PCP) and McyB_1_-(C^M213C^-A^R871C^) gave a 55-65% reduction in their adenylation activity as compared to the wild-type. Moreover, the reduction in this activity was reversed in both proteins by adding DTT (dithiothreitol) to reduce the disulfide bonds formed by the double Cys residues (Figure 4D and 4E). These results suggest that the interaction between the C and A domains at the top interface of McyA_2_-(C-A-PCP) and McyB_1_-(C-A) could be fixed by disulfide bond between the double Cys substitutions, which resulted in a reduction of the adenylation activity in both proteins. However, once the fixed conformation becomes flexible again by reducing the disulfide bonds, the adenylation activity could be restored to the original levels. Taken all together, we conclude that the dynamic interaction at the top interface between the C and A_sub_ domains is important for the activity of the A domain in the McyA_2_ and McyB_1_ modules during the catalytic cycle.

### Dynamic interaction between cognate C and A domains is functionally prevalent in NRPS elongation and termination modules

Since the dynamic interaction mediated by the “RGXR” motif enhances the activity of the McyA_2_ and McyB_1_ modules, we wondered if similar control mechanism also exists in other NRPS modules. We first analyzed the top interface between the A and C domains in all the reported NRPS structures corresponding to distinct stages of the catalysis. The interactions observed between the η10-α26 loop of the A_sub_ domain and the C domain in each respective structure mainly rely on non-bonded contacts, with notably fewer direct contacts (Figure S6). These results agree with the conclusion that the top interface between the C and A_sub_ domains of a NRPS module changes dynamically during the catalytic cycle. Next, we examined the sequence conservation of the “RGXR” motif in different NRPS modules, by aligning the A8 catalytic regions surrounding the “RGXR” motif from 99 NRPS modules of different organisms (Table S3). The results indicate that the “RGXR” motif is highly conserved in the elongation and termination modules of NRPSs, but it exhibits a very low sequence similarity in the initiation modules of NRPSs or the independent A domains (Figure 5A). These results suggest that the function in adenylation activity based on dynamic interaction between the “RGXR” motif and C domain may be specifically present in the elongation and termination modules of NRPSs.

**Figure 5.**
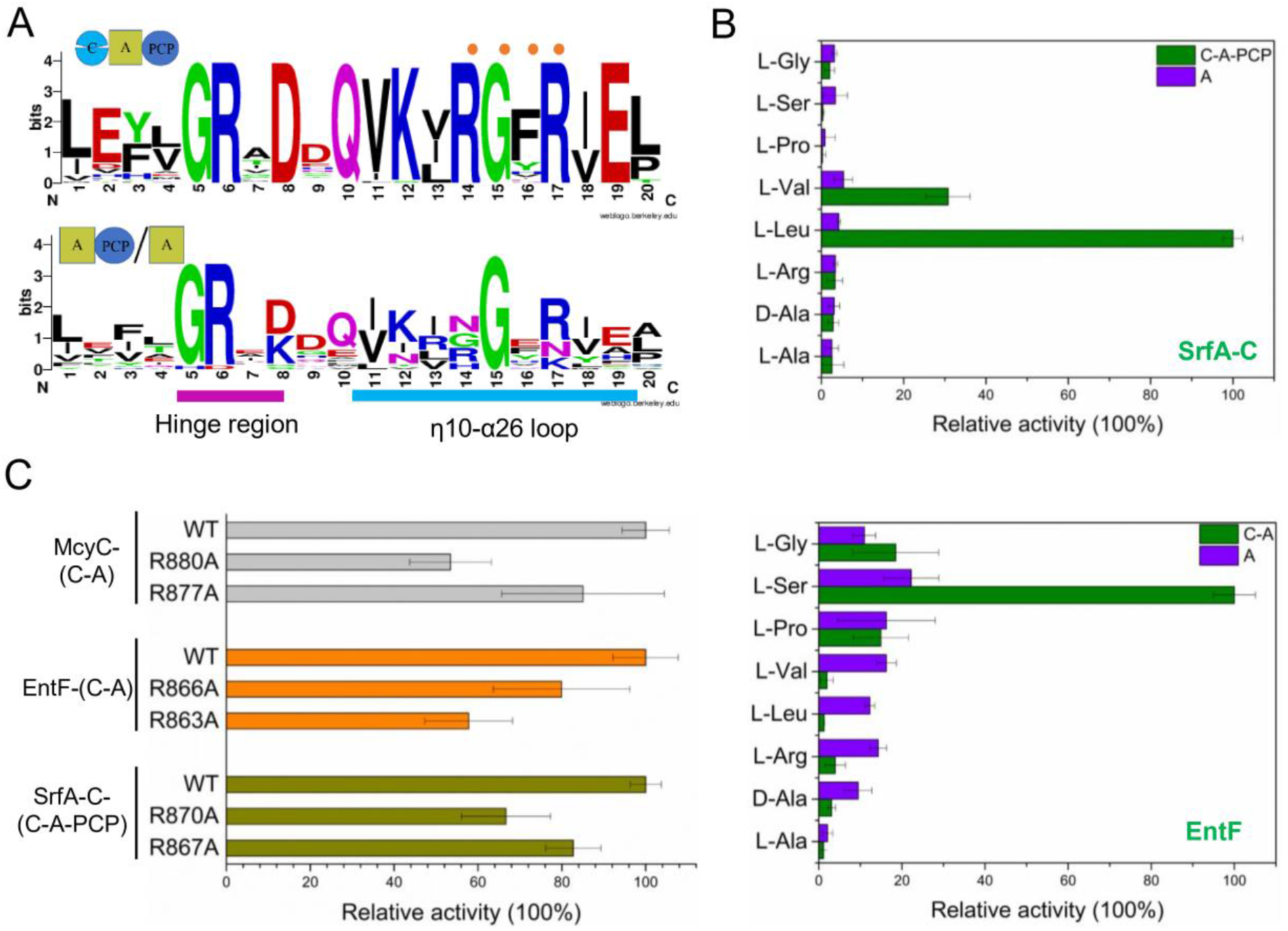
Functional characterization of dynamic interaction between A domain and C domains in different NRPS modules. (A) Sequence logo of the A8 catalytic motifs from different NRPS modules generated by WebLogo (http://weblogo.threeplusone.com) (Color scheme Chemistry). Heights of symbols within stack indicate the relative frequency of each amino acid residue at that position. The Logo at the upper panel was obtained through multiple alignment of the A8 catalytic motifs from 79 NRPS elongation or termination modules. The Logo at the bottom panel was obtained through multiple alignment of the A8 catalytic motifs from 21 NRPS initiation or standalone A domain modules. The conserved “hinge region” was indicated with a line in purple, and the η10-α26 loop with a line in blue. The “RGXR” motif is indicated by orange dots. (B) In vitro adenylation activities of different truncated polypeptides derived from SrfA-C (upper panel) and EntF (bottom panel). (C) The adenylation activation profiles of McyC-(C-A), SrfA-C-(C-A-PCP) and EntF-(C-A) toward their respective specific substrates (L-Arg for McyC-(C-A), L-Leu for SrfA-C-(C-A-PCP), L-Ser for EntF-(C-A)) with or without point mutations. The adenylation activities were determined using a continuous hydroxylamine release assay. The OD_650_ value of the WT protein is set to 100%. Error bars indicate standard deviation based on the mean of three experimental replicates.

To validate this possibility, three representative NRPS modules, McyC, SrfA-C, EntF, were selected for further investigation. SrfA-C is the termination module for surfactin synthesis in *Bacillus subtilis* ATCC 21332, and it catalyzes the addition of L-leucine to the hexapeptide precursor and releases the end product by cyclization (Tanovic *et al*., 2008). EntF is the termination module responsible for L-serine activation during the biosynthesis of enterobactin in *E. coli* (Miller *et al*, 2016b). Comparison of the adenylation activity of the C-A-PCP tridomain or C-A didomain recombinant polypeptides of SrfA-C or EntF with their corresponding ones with the A domains alone revealed that the presences of the C domain in the SrfA-C and the EntF modules also enhanced their respective adenylation activity (L-Leu to SrfA-C, L-Ser to EntF) (Figure 5B). These results are thus similar to those observed with the McyA_2_, the McyB_1_ and the McyC modules (Figure 1). When the conserved two Arg residues in the “RGXR” motif of McyC-(C-A), SrfA-C-(C-A-PCP) and EntF-(C-A) were replaced, respectively, by Ala, the variants all exhibited a reduction in various degrees in the adenylation activity (Figure 5C). These results suggested that, similar to the McyA_2_ and the McyB_1_ modules, the dynamic interaction mediated by the “RGXR” motif between the C and the A_sub_ domains present in McyC, SrfA-C, EntF modules and is also important for the adenylation activity of these modules. Therefore, the function of the dynamic interaction between the adjacent C and A_sub_ domains mediated by the “RGXR” motif is prevalent in different NRPS elongation and termination modules.

## Discussion

A NRPS module contains multiple domains working together for the incorporation of a single residue into the growing peptide chain in an orderly catalytic cycle. The highly dynamic architecture of these domains is considered to be the key for efficient catalytic reactions (Bonhomme *et al*, 2021; Izore & Cryle, 2018; Patel *et al*., 2023; Winn *et al*, 2016). Due to the dynamic nature of such multidomain proteins, it is difficult to capture the structures of the intermediate conformations of a complete NRPS module, which limits our understanding on the structural basis of the catalytic mechanisms. In this study, we obtained two crystal structures of the NRPS involved in MC biosynthesis, one for a truncated module McyB_1_-(C-A), and another for a complete elongation module McyA_2_-(C-A-PCP) (Figure 2). Structural analysis and comparison with the seven reported NRPS structures that all contain both a C and an A domains show that the structures of McyA_2_-(C-A-PCP) and McyB_1_-(C-A) reported here represent two newly identified conformations of NRPS in a catalytic cycle (Figure 2 and 3, Figure S4) (Drake *et al*., 2016; Kreitler *et al*., 2019; Reimer *et al*., 2019; Tan *et al*, 2020; Tanovic *et al*., 2008). For McyB_1_-(C-A), the absence of the PCP domain in the module prevents the transfer of the activated substrate from the A domain to the C domain, resulting in a conformation locked at the adenylation step of the catalytic cycle (Figure 2A and 3A). A similar C-A bidomain structure of a NRPS module involved in teixobactin production was previously reported (Txo2_C_1_-A_1_) (Tan *et al*., 2020); however, the substrate and AMP were not present in the pocket of the active site in this structure, even though the A_sub_ domain is closed upon the active site. Thus, the reported structure of Txo2_C_1_-A_1_ is not a perfect one to mimic the adenylation state of the corresponding NRPS module. The presence of the substrate in the structure of McyB_1_-(C-A) reported here may better capture the structural features of the adenylation state during the catalytic cycle. For McyA_2_-(C-A-PCP), before setting up for crystallization, the PPant transferase (Sfp enzyme) was used to activate the phosphopantetheinyl (PPant) arm of the PCP domain (Quadri *et al*, 1998). Consequently, the C domain in the McyA_2_-(C-A-PCP) structure displays a conformation in which the PPant arm of the PCP domain is docked into the cleft of the C domain, a situation similar to that for those structures of modules treated by a similar process, such as AB3403 (Drake *et al*., 2016), ObiF1 (Kreitler *et al*., 2019) and LgrA (Reimer *et al*., 2019). However, unlike the A domains of these three proteins, which are positioned in the “adenylation (closed)” conformation, the A domain in our structure is in a “pre-adenylation” conformation where the substrate and the A_sub_ domain were not yet positioned for the adenylation reaction to occur (Figure 3C and 3D). This pre-adenylation structure is thus characteristic of an intermediate catalytic conformation of NRPS.

All complete module structures of NRPSs previously reported correspond to three different C–A domain conformation combinations, representing three steps in the catalytic cycle. These findings indicate that the C and A domains constitute a catalytic platform, upon which the other domains can assemble and move along (Drake *et al*., 2016; Izore & Cryle, 2018; Patel *et al*., 2023; Tanovic *et al*., 2008). The structures of McyA_2_-(C-A-PCP) and McyB_1_-(C-A) obtained in this study provide two additional distinct conformations for C–A didomains during catalysis. In all these structures, the bottom interfaces for interaction between the C domain and the A_core_ domain is stable, and the dynamic changes occur mainly at the level of the top interfaces between the A_sub_ and the C domains, caused by rotation of the A_sub_ domain during catalysis (Figure 4A) (Drake *et al*., 2016; Kreitler *et al*., 2019; Tanovic *et al*., 2008). Direct contact interactions at the top interfaces are not obvious in either the McyB_1_-(C-A) structure or other structures (Figure 4C and Figure S6); however, we found that salt bridges and hydrogen bonds are formed between the η10-α26 loop of the A_sub_ domain and the C domain in the McyA_2_-(C-A-PCP) structure (Figure 4). This is the first time that interactions based on such strong direct contacts have been observed at the flexible top interface between the C domain and the A domain of a NRPS module. These observations suggest that certain direct interaction between the A_sub_ and C domains may happen at the intermediate stages during in the NRPS catalytic cycle.

Beyond expanding the interaction of C-A didomains in NRPS modules during the catalytic cycle, our studies provided a detailed functional characterization on the newly discovered interactions at this region. Previous studies found that deletion of the C domain alters the adenylation activity for the McyB_1_ and McyC NRPS modules required for MC synthesis and the module for sulfazecin production (Li *et al*., 2017; Meyer *et al*., 2016). Our data reported here not only provided structural insight into the participation of the C domains in the adenylation reaction, but also indicated that this effect is prevalent in elongation and termination modules of NRPSs (Figure 1 and Figure 5). Our studies demonstrated that in these modules, dynamic interaction at the top interfaces between the C and A_sub_ domains is necessary for optimal adenylation activity. Based on the direct interaction at this region observed in the structure of McyA_2_-(C-A-PCP) and the results of sequence alignment, key residues of the “RGXR” motifs responsible for formation of salt bridges and hydrogen bonds in the η10-α26 loop of the A_sub_ domains are highly conserved in different elongation and termination modules of NRPSs (Figure 5A). Moreover, disrupting the sidechain of the two conserved Arg residues in the “RGXR” motif resulted in a reduction of the adenylation activity in different NRPS modules tested (Figure 4 and 5). These results suggest that the function of the newly discovered interaction region is widespread in NRPSs and important for catalysis.

A significant conformational rearrangement of the A domain, in particular the A_sub_ domain, is the key for an efficient catalysis during the reaction. The rotation of the A_sub_ domain is accompanied by a change in the angle of the two conserved “hinge” residues (R and D) at the conserved A8 motif required for connecting the A_core_ and A_sub_ domains (Figure 5A) (Reger *et al*, 2007; Wu *et al*, 2009). Since the conserved η10-α26 loop is also a part of the A8 catalytic motif in the NRPS elongation and termination modules, just after the “hinge” residues (Figure 5A), it is possible that the conserved “RGXR” motif in the η10-α26 loop plays an important role in assisting the rotation of the A_sub_ domain and in coordinating the catalytic reaction of the adjacent A and C domains during catalysis. Based on previous studies and our findings, we propose a model on the control mechanism of the adenylation activity in a NRPS module mediated by dynamic interaction between the two domains (Figure 6). When the substrate of a NRPS module binds at the pocket of the A_core_ domain, it triggers a change at a right angle of the “hinge” residues located between the two subdomains. Subsequently, the dynamic interaction between the “RXGR” motif and the C domain guides the rotational direction of the A_sub_ domain, enabling a more efficient state transition and a high adenylation efficiency of the A domain. When the dynamic interaction between the “RXGR” motif and the C domain was disrupted, the A_sub_ domain rotates in a random fashion, thereby resulting in a low adenylation efficiency of the A domain in a NRPS module.

**Figure 6.**
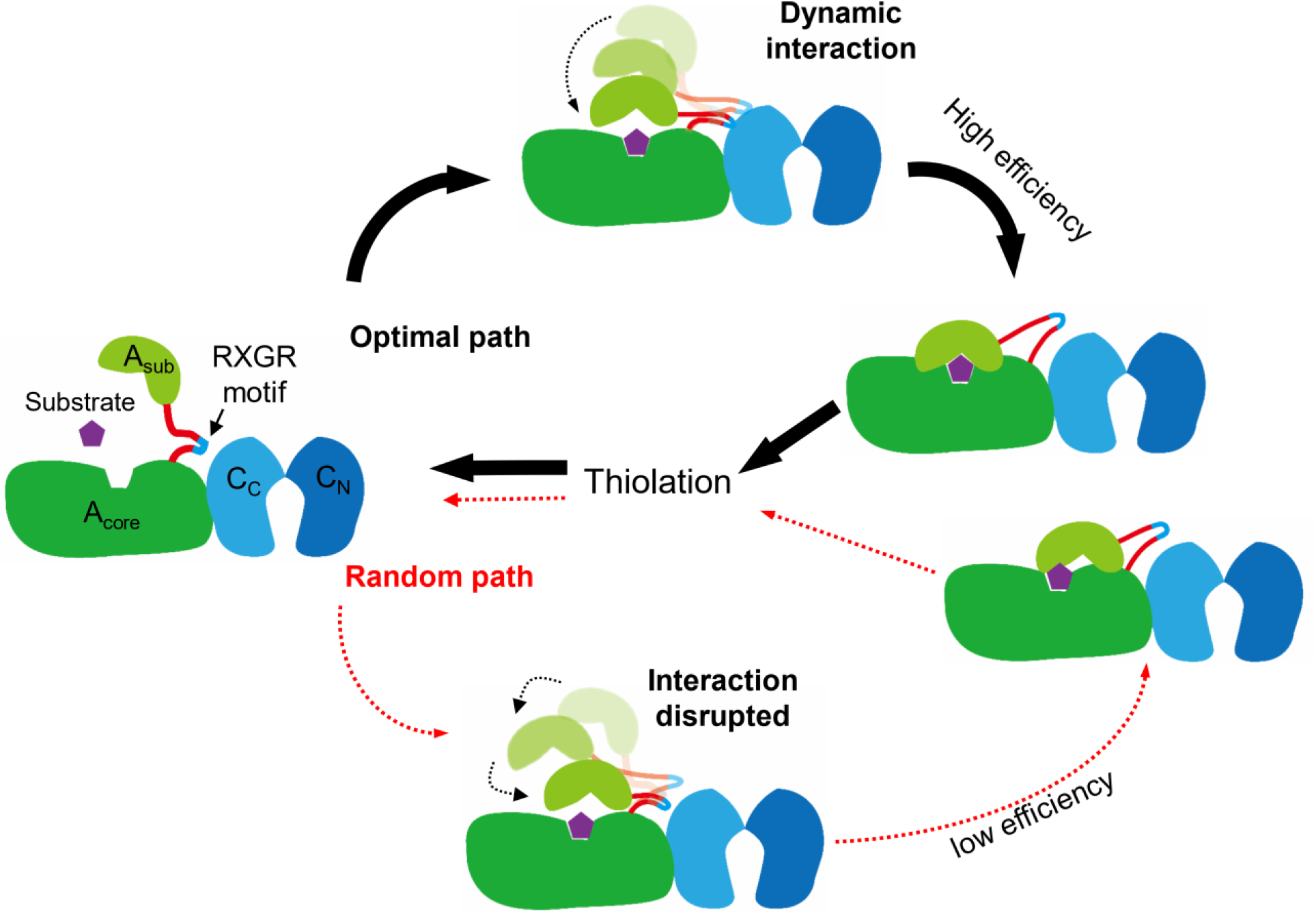
A model for the control mechanism of the adenylation activity in a NRPS module, mediated by dynamic interaction between the A and C domains. Binding of the substrate in a NRPS module triggers a change of the “hinge” region (red line) located at the linker between the A_sub_ domain and the A_core_ domain. Subsequently, dynamic interaction between the “RXGR” motif which after the hinge region and the C domain guide the rotation of the A_sub_ domain in a direction that allows an efficient state transition (from the open conformation to adenylation/closed conformation) of the A domain. In this case, the NRPS module shows a highly efficient adenylation activity (optimal path, upper part, with thick arrows). When the interaction between the “RXGR” motif and the C domain was disrupted, the rotation of the A_sub_ domain occurs rather randomly with ill fitted state transition or conformational changes, and thereby, the A domain in the NRPS module shows adenylation activity with poor efficiency.

Certain details in this model still need to be understood, in particular more intermediate structural snapshots are required for a comprehensive understanding of the catalytic cycle. Nevertheless, the model provides a framework for the underlying control mechanism involved in the catalytic activity in the presence of a substrate in NRPS modules. NRPSs are attractive targets for bioengineering for peptide products in the era of synthetic biology. Understanding the control mechanism for catalysis will facilitate bioengineering of NRPSs to design and generate new peptidic products.

## Methods

### Plasmids construction

The plasmids, used for protein expression and purification, were constructed based on the pET28a expression vector, which contains the sequence encoding a 6XHis tag. DNA fragments encoding the single A domain, the C-A and A-PCP didomains, or the C-A-PCP tri-domain polypeptides of McyA_2_, McyB_1_ and McyC were PCR amplified, respectively, from the *Microcystis* genomic DNA (GenBank: AM778952.1). DNA fragments encoding the single A domain, or C-A-PCP tri-domain of EntF were PCR amplified, respectively, from *E. coli* JM109 (The primers were designed according to the K-12 genome sequence, GenBank: NC_000913.3). DNA fragments encoding the single A domain or C-A didomain of SrfA-C were amplified, respectively, by PCR from genomic DNA of the surfactin producer strain *B. subtilis* ATCC 21332. The amplified DNA fragments were cloned into pET28a, transformed into *E. coli* DH5α, and confirmed by sequencing. To construct the plasmids containing genes with point mutation(s), quick-change mutagenesis was carried out using the plasmid containing the corresponding WT genes as template. Plasmids, and oligonucleotides used in this study are listed in Table S1 and S2, respectively.

### Protein expression and purification

The plasmids for recombinant protein expression were transformed into *E. coli* BL21 (DE3) and the strains were grown in Luria-Bertani (LB) medium containing kanamycin (50 mg/ml) with shaking at 37°C until OD_600_ of 0.4-0.6. Protein expression was induced overnight with addition of 0.5 mM isopropyl-β-d-thiogalactoside (IPTG) at 16°C. Cells were collected by centrifugation at 9000 × g for 10 min. Selenomethionine (SeMet)-labeled proteins were overexpressed in *E. coli* BL21 (DE3) (Novagen). Transformed cells were first cultured in LB medium at 37°C overnight, then harvested and washed twice with M9 medium (Walden, 2010). Then cells were cultured in SeMet medium (M9 medium with 50 mg·L^−1^ SeMet and other essential amino acids) to OD_600_ of 0.6–0.8.

The native and SeMet labeled proteins was produced as described previously, with minor modifications.(Zeng *et al*, 2023) Cell pellets were resuspended in lysis buffer (200 mM NaCl, 10% glycerin, 20 mM Tris-HCl pH 8.0, 0.2 mM TCEP). Cells were lysed using French press (JINBO,Inc), followed by centrifugation at 12000 × g for 40 min at 4°C. The initial purification was achieved with a Ni-NTA column. The supernatant was mixed with 10% (v/v) Ni-NTA agarose beads (Qiagen) preequilibrated with the lysis buffer for 40 min at 4°C. The beads were then collected by filtration and washed with the lysis buffer and the washing buffer (200 mM NaCl, 10% glycerol, 20 mM Tris-HCl pH 8.0, 0.2 mM TCEP, 20 mM imidazole). Protein was eluted using elution buffer (200 mM NaCl, 10% glycerol, 20 mM Tris-HCl pH 8.0, 0.2 mM TCEP, 250 mM imidazole). The eluted samples were concentrated (Amicon Ultracel-10K, Millipore) and further purified by size-exclusion chromatography (SEC) with a HiLoad 26/600 Superdex 200 column (GE Healthcare). The column was equilibrated with the SEC buffer (200 mM NaCl, 10% glycerol, 20 mM Tris-HCl pH 8.0, 0.2 mM TCEP). The purity of each fraction was determined by SDS-PAGE. Fractions containing the pure recombinant protein were pooled, concentrated and stored at −80°C for further use.

### Crystallization

Before setting up for crystallization, purified native proteins or SeMet-labelled McyA_2_-(C-A-PCP) were phosphopantetheinylated by incubation with the phosphopantetheinyl transferase Sfp (10 nM), in the presence of 12.5 mM MgCl_2_ and 1 mM CoA for 60 min at 20°C. Then, the protein sample was passed over a HiLoad 26/600 Superdex 200 column (GE Healthcare) pre-equilibrated with the equilibration buffer (100 mM NaCl, 20 mM Tris-HCl pH 8.0, 0.2 mM TCEP) to remove the Sfp protein. The eluted holo-McyA_2_-(C-A-PCP) protein was pooled and concentrated to 15 mg/ml by ultrafiltration (Amicon Ultracel-10K, Millipore), then, mixed with 2 mM ATP, 2 mM L-Ala and 3 mM MgCl_2_ at 4°C for 1 h. The crystallization conditions were initially identified from a sparse matrix screen at 20°C. Final crystals for SeMet-labelled holo-McyA_2_-(C-A-PCP) were obtained against the reservoir solution containing 0.18 M lithium sulfate monohydrate,0.1 M Tris-Cl pH 8.0,17% w/v polyethylene glycol 3350 using the hanging drop vapor diffusion method at 20°C, with a drop of 1 μL protein solution mixed with 1 μL reservoir.

For McyB_1_-(C-A), the purified native protein was first concentrated to 8 mg/ml by ultrafiltration (Amicon Ultracel-10K, Millipore), then mixed with 2 mM ATP, 2 mM L-Leu and 3 mM MgCl_2_ at 4°C for 1 h. The initial crystallization conditions for the McyB_1_-(C-A) were identified from a sparse matrix screen at 20°C. Final high-quality crystals were grown against the reservoir solution containing 3.5 M sodium formate, 5% glycerol, 80 mM sodium chloride, and 50 mM Tris-HCl pH 8.0 using hanging drop vapor diffusion at 20°C, with a drop of 1 μL protein solution mixed with 1 μL reservoir.

### Structure determination of McyA_2_-(C-A-PCP) and McyB_1_-(C-A)

SeMet-labelled crystals of holo-McyA_2_-(C-A-PCP) were flash-cooled with 30% glycerol as cryoprotectant in liquid nitrogen. X-ray diffraction data was collected at 100 K on an ADSC Q315r CCD detector at beamline BL19U of Shanghai Synchrotron Radiation Facility (SSRF). Diffraction data was indexed, integrated and scaled using *HKL2000 (Otwinowski & Minor, 1997)* in space group *P3121*. The structure was solved by molecular replacement with *Molrep* in CCP4i (Winn *et al*, 2011). Initially, we solved the structure of the C domain using the AlphaFold (Jumper *et al*, 2021) predicted model of McyA_2_ (residue I1267-S1710) as an initial search model. Afterwards, via fixing the solution of the C domain, the MR was performed using the A domain (residue E1711-S2224, predicted by AlphaFold) of McyA_2_ as a subsequent search model. Finally, the PCP domain was solved by MR using the model of AB3403 (PDB:4ZXH, residues A971-E1045) via fixing the C and A domains. The maximum-likelihood method implemented in *REFMAC5* (Murshudov et al, 2011) as part of the *CCP4* program suite (Winn *et al*., 2011) and *Coot* (Emsley & Cowtan, 2004) were used interactively to build the linkage among different domains manually, and the structure was automatically refined in the program Phenix (Adams *et al*, 2002). The ligands and waters were manually built in the *F_O_-F_C_* maps in Coot to produce the final model. The ATP molecule of McyA_2_-(C-A-PCP) was manually adjusted in Coot and automatically refined in REFMAC5 of CCP4. The final model was evaluated with *PROCHECK* (Laskowski et al, 1993) and *MolProbity* (Chen et al, 2010).

Crystals of McyB_1_-(C-A) were flash-cooled with 20% glycerol as cryoprotectant in liquid nitrogen. In total, 1157 images of diffraction data were collected using a Rigaku RUH-3R rotating copper-anode source equipped with a PILATUS 200K detector (DECTRIS, Switzerland) at 100 K. Diffraction data was indexed, merged, and scaled using *iMOSFLM* (Battye et al, 2011) in space group *I 41* (2.70 Å). Structure determination was performed in *PHENIX* using molecular replacement and the structure of AB3403 (PDB: 4ZXH, residues M1-E964) was used as the search model. The subsequent refinement process is similar to holo-McyA_2_-(C-A-PCP). Crystallographic parameters and data-collection statistics for both structures are listed in Table 1.

**Table 1.** Crystallographic Data Statistics.

|  | McyB<sub>1</sub>-(C-A) (Native)* | McyA<sub>2</sub>-(C-A-PCP) (SeMet)* |
| --- | --- | --- |
| **Data Collection** |  |  |
| Beamline | RIGAKU MICROMAX-007 | SSRF- BL19U1 |
| Wavelength (Å) | 1.5418 | 0.9785 |
| Space group | P3121 | I41 |
| Unit Cell Dimensions |  |  |
| a, b, c (Å) | 152.53, 152.53, 135.74 | 174.01, 174.01, 78.03 |
| α, β, γ (°) | 90.00, 90.00, 120.00 | 90.00, 90.00, 90.00 |
| Resolution range (Å) | 13.15-2.70 (2.79-2.70) | 43.50-2.45 (2.54-2.45) |
| R<sub>meas</sub> | 0.129 (1.549) | 0.118 (0.733) |
| Redundancy | 8.0 (6.1) | 11.0 (5.9) |
| Completeness (%) | 99.9 (98.9) | 99.9 (99.9) |
| CC<sub>1/2</sub> | 0.997 (0.679) | 0.846 (0.721) |
| I/σ(I) | 17.5 (2.2) | 19.1 (2.3) |
| **Refinement** |  |  |
| Resolution range (Å) | 13.15-2.70 | 43.50-2.45 |
| Average B factor (Å<sup>2</sup>) | 69.00 | 81.00 |
| No. of reflections | 47370 | 40803 |
| Completeness | 98.9 | 99.9 |
| R<sub>work</sub> /R<sub>free</sub> | 0.213/0.269 | 0.232/0.305 |
| No. of protein (residues) | 910 | 1038 |
| No. of ligand (atoms) | 31 | 51 |
| RMSD bond lengths (Å) | 0.008 | 0.006 |
| RMSD bond angles (°) | 1.701 | 1.404 |
| **Ramachandran Statistics** |  |  |
| Favored (%) | 91.0 | 89.0 |
| Allowed (%) | 8.0 | 10.0 |
| Outliers (%) | 1.0 | 1.0 |
| **PDB code** | **8HLK** | **8JBR** |
| *Statistics for the highest-resolution shell are shown in parentheses. |  |  |

### Continuous Hydroxylamine Release Assay

The adenylation activity of A domains in different proteins was determinated by the continuous hydroxylamine release assay (Figure S2). This method is similar to the hydroxylamine release assay described previously, with minor modifications (Duckworth *et al*, 2016). Briefly, 300 µL freshly prepared solution A (0.04 U inorganic phosphatase, 150 mM hydroxylamine, 5 mM ATP, 5 mM MgCl_2_,100 mM Tris-HCl pH 8.0) was first mixed with 6 µl 100 mM amino acid solution at 25°C for 5 min. The adenylation reaction was initiated by addition of 6 µl (1 µM / L) protein in 1.5-ml tubes at 25°C for 10 min. Then, 300 µl dyeing solution (Phosphate colorimetric assay kit, Sigma-Aldrich) was transferred into the reaction tubes and incubated at 25°C for 30 min. After incubation, 200 µl reaction solution was transferred into a 96-well plate and the absorbance at 650 nm measured by using Spectramax M5e.

### Bioinformatic analysis

The sequence identity or similarity of NPRS homologs with C-A didomains was calculated using Clustal Omega software. Multiple sequence alignment was presented by ESPript 3.0 with 5 representative sequences of C-A didomains from *Microcystis aeruginosa* PCC 7806, *Escherichia coli* JM109 and *Bacillus subtilis* ATCC 21332. The sequence logo of the A8 catalytic motif in NRPS elongation or termination modules are made by the WebLogo server with 79 representative modules (Table S3). The sequence logo of the A8 catalytic motif in initiation module or standalone A domain of different NRPSs are made by the WebLogo server with 21 representative NRPS module (Table S3). These NRPSs are from different bacteria, including *Microcystis aeruginosa* PCC 7806, *Escherichia coli* JM109, *Bacillus subtilis* ATCC 21332, *Acinetobacter baumannii* ATCC 17978, *Brevibacillus parabrevis* ATCC 8185, *Burkholderia diffusa*, *Streptomyces atroolivaceus*, *Pseudomonas protegens* Pf-5, *Pantoea agglomerans*, *Bacillus velezensis* YAU B9601-Y2, *Streptomyces rapamycinicus* NRRL 5491, *Amycolatopsis rubida*, *Brevibacillus* sp. CF 112, *Brevibacillus* sp. BC 25, *Amycolatopsis orientalis*, *Bacillus licheniformis*, *Nodularia spumigena* UHCC 0039, *Bacillus brevis* ATCC 999, *Streptomyces roseosporus* NRRL 11379, *Mycobacterium smegmatis* ATCC 700084, *Pseudomonas aeruginosa* PAO1 and *Streptomyces halstedii*.

### Data availability section

The accession numbers for the structures reported in this paper are PDB: 8HLK, 8JBR. All figures showing a structure were prepared with PyMOL (https://pymol.org/2/).

## Acknowledgments

The authors are thankful to staff at Shanghai Synchrotron Radiation Facility for help in data collection. This research was supported by the National Key R&D Program of China (project number 2018YFA0903100) and the Youth Innovation Promotion Association CAS.

## Author contributions

Conceptualization, CC.Z., CZ.Z. Y.C. and X.Z.; Methodology, YJ. P. and YL.J.; Investigation, YJ. P; Formal Analysis, YJ. P., X.Z. and YL. J.; Visualization, YJ.P. and X.Z.; Writing – Original Draft: X.Z and YJ.P., Writing – Review & Editing: X.Z., YL.J., CZ.Z., Y.C., W.M. and CC.Z.; Funding Acquisition: CC.Z. and X.Z.

## References

Adams PD, Grosse-Kunstleve RW, Hung L-W, Ioerger TR, McCoy AJ, Moriarty NW, Read RJ, Sacchettini JC, Sauter NK, Terwilliger TC (2002) PHENIX: building new software for automated crystallographic structure determination. Acta Crystallographica Section D 58: 1948–1954

Battye TGG, Kontogiannis L, Johnson O, Powell HR, Leslie AGW (2011) iMOSFLM: a new graphical interface for diffraction-image processing with MOSFLM. Acta Crystallographica Section D 67: 271–281

Bloudoff K, Rodionov D, Schmeing TM (2013) Crystal Structures of the First Condensation Domain of CDA Synthetase Suggest Conformational Changes during the Synthetic Cycle of Nonribosomal Peptide Synthetases. Journal of Molecular Biology 425: 3137–3150

Bonhomme S, Dessen A, Macheboeuf P (2021) The inherent flexibility of type I non-ribosomal peptide synthetase multienzymes drives their catalytic activities. Open Biology 11: 200386

Botes DP, Tuinman AA, Wessels PL, Viljoen CC, Hammond SJ (1984) The structure of cyanoginosin-LA, a cyclic heptapeptide toxin from the cyanobacterium Microcystis aeruginosa. Journal of the Chemical Society Perkin Transactions 1 1

Chen VB, Arendall WB, III, Headd JJ, Keedy DA, Immormino RM, Kapral GJ, Murray LW, Richardson JS, Richardson DC (2010) MolProbity: all-atom structure validation for macromolecular crystallography. Acta Crystallographica Section D 66: 12–21

Conti E, Stachelhaus T, Marahiel MA, Brick P (1997) Structural basis for the activation of phenylalanine in the non-ribosomal biosynthesis of gramicidin S. Embo J 16: 4174–4183

Crosby J, Crump MP (2012) The structural role of the carrier protein – active controller or passive carrier. Nat Prod Rep 29: 1111–1137

Dittmann E, Neilan BA, Erhard M, Döhren Hv, Börner T (1997) Insertional mutagenesis of a peptide synthetase gene that is responsible for hepatotoxin production in the cyanobacterium Microcystis aeruginosa PCC 7806. Molecular Microbiology 26: 779–787

Drake EJ, Miller BR, Shi C, Tarrasch JT, Sundlov JA, Allen CL, Skiniotis G, Aldrich CC, Gulick AM (2016) Structures of two distinct conformations of holo-non-ribosomal peptide synthetases. Nature 529: 235–238

Drake EJ, Nicolai DA, Gulick AM (2006) Structure of the EntB multidomain nonribosomal peptide synthetase and functional analysis of its interaction with the EntE adenylation domain. Chem Biol 13: 409–419

Duckworth BP, Wilson DJ, Aldrich CC (2016) Measurement of Nonribosomal Peptide Synthetase Adenylation Domain Activity Using a Continuous Hydroxylamine Release Assay. In: Nonribosomal Peptide and Polyketide Biosynthesis: Methods and Protocols, Evans B.S. (ed.) pp. 53-61. Springer New York: New York, NY

Duy TN, Lam PKS, Shaw GR, Connell DW (2000) Toxicology and risk assessment of freshwater cyanobacterial (blue-green algal) toxins in water. Reviews of Environmental Contamination & Toxicology 163: 113

Emsley P, Cowtan K (2004) Coot: model-building tools for molecular graphics. Acta Crystallographica Section D 60: 2126–2132

Felnagle EA, Jackson EE, Chan YA, Podevels AM, Berti AD, McMahon MD, Thomas MG (2008) Nonribosomal Peptide Synthetases Involved in the Production of Medically Relevant Natural Products. Molecular Pharmaceutics 5: 191–211

Fischbach MA, Walsh CT (2006) Assembly-Line Enzymology for Polyketide and Nonribosomal Peptide Antibiotics: Logic, Machinery, and Mechanisms. Chemical Reviews 106: 3468–3496

Frueh DP, Arthanari H, Koglin A, Vosburg DA, Bennett AE, Walsh CT, Wagner G (2008) Dynamic thiolation–thioesterase structure of a non-ribosomal peptide synthetase. Nature 454: 903–906

Gulick AM (2009) Conformational dynamics in the Acyl-CoA synthetases, adenylation domains of non-ribosomal peptide synthetases, and firefly luciferase. Acs Chem Biol 4: 811–827

Izore T, Cryle MJ (2018) The many faces and important roles of protein-protein interactions during non-ribosomal peptide synthesis. Nat Prod Rep 35: 1120–1139

Jumper J, Evans R, Pritzel A, Green T, Figurnov M, Ronneberger O, Tunyasuvunakool K, Bates R, Žídek A, Potapenko A et al (2021) Highly accurate protein structure prediction with AlphaFold. Nature 596: 583–589

Kaniusaite M, Tailhades J, Marschall EA, Goode RJA, Cryle MJ (2019) A proof-reading mechanism for non-proteinogenic amino acid incorporation into glycopeptide antibiotics. Chemical Science 10

Keating TA, Marshall CG, Walsh CT, Keating AE (2002) The structure of VibH represents nonribosomal peptide synthetase condensation, cyclization and epimerization domains. Nature Structural Biology 9: 522–526

Koglin A, Mofid MR, Löhr F, Schäfer B, Rogov VV, Blum M-M, Mittag T, Marahiel MA, Bernhard F, Dötsch V (2006) Conformational Switches Modulate Protein Interactions in Peptide Antibiotic Synthetases. Science 312: 273–276

Koglin A, Walsh CT (2009) Structural insights into nonribosomal peptide enzymatic assembly lines. Nat Prod Rep 26: 987–1000

Konz D, Marahiel MA (1999) How do peptide synthetases generate structural diversity? Chem Biol 6: R39

Kreitler DF, Gemmell EM, Schaffer JE, Wencewicz TA, Gulick AM (2019) The structural basis of N-acyl-α-amino-β-lactone formation catalyzed by a nonribosomal peptide synthetase. Nature Communications 10: 3432

Laskowski RA, Macarthur MW, Moss DS, Thornton JM (1993) PROCHECK: a program to check the stereochemical quality of protein structures. Journal of Applied Crystallography 26

Lawson DM, Derewenda U, Serre L, Ferri S, Szittner R, Wei Y, Meighen EA, Derewenda ZS (1994) Structure of a Myristoyl-ACP-Specific Thioesterase from Vibrio harveyi. Biochemistry-Us 33: 9382–9388

Li R, Oliver RA, Townsend CA (2017) Identification and Characterization of the Sulfazecin Monobactam Biosynthetic Gene Cluster. Cell Chemical Biology

Massey IY, Al Osman M, Yang F (2022) An overview on cyanobacterial blooms and toxins production: their occurrence and influencing factors. Toxin Rev 41: 326–346

Meiβner K, Dittmann E, B?Rner T (1996) Toxic and non-toxic strains of the cyanobacterium Microcystis aeruginosacontain sequences homologous to peptide synthetase genes. FEMS Microbiol Lett 135: 295-303. Fems Microbiology Letters 135: 295–303

Meyer S, Kehr JC, Mainz A, Dehm D, Dittmann E (2016) Biochemical Dissection of the Natural Diversification of Microcystin Provides Lessons for Synthetic Biology of NRPS. Cell Chemical Biology 23: 462–471

Miller BR, Drake EJ, Shi C, Aldrich CC, Gulick AM (2016a) Structures of a Nonribosomal Peptide Synthetase Module Bound to MbtH-like Proteins Support a Highly Dynamic Domain Architecture. J Biol Chem 291: 22559–22571

Miller BR, Drake EJ, Shi C, Aldrich CC, Gulick AM (2016b) Structures of a Nonribosomal Peptide Synthetase Module Bound to MbtH-like Proteins Support a Highly Dynamic Domain Architecture. J Biol Chem: jbc.M116.746297

Mitchell CA, Shi C, Aldrich CC, Gulick AM (2012) Structure of PA1221, a Nonribosomal Peptide Synthetase Containing Adenylation and Peptidyl Carrier Protein Domains. Biochemistry-Us 51: 3252–3263

Murshudov GN, Skubak P, Lebedev AA, Pannu NS, Steiner RA, Nicholls RA, Winn MD, Long F, Vagin AA (2011) REFMAC5 for the refinement of macromolecular crystal structures. Acta Crystallographica Section D 67: 355–367

Otwinowski Z, Minor W (1997) Processing of X-ray diffraction data collected in oscillation mode. Method Enzymol 276: 307–326

Patel KD, MacDonald MR, Ahmed SF, Singh J, Gulick AM (2023) Structural advances toward understanding the catalytic activity and conformational dynamics of modular nonribosomal peptide synthetases. Nat Prod Rep

Quadri LEN, Weinreb PH, Lei M, Nakano MM, Zuber P, Walsh CT (1998) Characterization of Sfp, a Bacillus subtilis phosphopantetheinyl transferase for peptidyl carrier protein domains in peptide synthetases. Biochemistry-Us 37: 1585–1595

Reger AS, Carney JM, Gulick AM (2007) Biochemical and crystallographic analysis of substrate binding and conformational changes in Acetyl-CoA synthetase. Biochemistry-Us 46: 6536–6546

Reger AS, Wu R, Dunaway-Mariano D, Gulick AM (2008) Structural Characterization of a 140° Domain Movement in the Two-Step Reaction Catalyzed by 4-Chlorobenzoate:CoA Ligase. Biochemistry-Us 47: 8016–8025

Reimer JM, Aloise MN, Harrison PM, Schmeing TM (2016) Synthetic cycle of the initiation module of a formylating nonribosomal peptide synthetase. Nature 529: 239–U305

Reimer JM, Eivaskhani M, Harb I, Guarné A, Weigt M, Schmeing TM (2019) Structures of a dimodular nonribosomal peptide synthetase reveal conformational flexibility. Science 366: eaaw4388

Rinehart KL, Namikoshi M, Choi BW (1994) Structure and biosynthesis of toxins from blue-green-algae (Cyanobacteria). Journal of Applied Phycology 6: 159–176

Samel SA, Schoenafinger G, Knappe TA, Marahiel MA, Essen L-O (2007) Structural and Functional Insights into a Peptide Bond-Forming Bidomain from a Nonribosomal Peptide Synthetase. Structure 15: 781–792

Strieker M, Tanovic A, Marahiel MA (2010) Nonribosomal peptide synthetases: structures and dynamics. Curr Opin Struc Biol 20: 234–240

Süssmuth RD, Mainz A (2017) Nonribosomal Peptide Synthesis—Principles and Prospects. Angewandte Chemie International Edition 56: 3770–3821

Tan K, Zhou M, Jedrzejczak RP, Wu R, Joachimiak A (2020) Structures of Teixobactin-producing Nonribosomal Peptide Synthetase Condensation and Adenylation Domains. Current Research in Structural Biology 2: 14–24

Tan XF, Dai YN, Zhou K, Jiang YL, Ren YM, Chen YX, Zhou CZ (2015) Structure of the adenylation-peptidyl carrier protein didomain of the Microcystis aeruginosa microcystin synthetase McyG. Acta Crystallogr D 71: 873–881

Tanovic A, Samel SA, Essen L-O, Marahiel MA (2008) Crystal Structure of the Termination Module of a Nonribosomal Peptide Synthetase. Science 321: 659–663

Tillett D, Dittmann E, Erhard M, von Döhren H, Börner T, Neilan BA (2000) Structural organization of microcystin biosynthesis in Microcystis aeruginosa PCC7806: an integrated peptide–polyketide synthetase system. Chem Biol 7: 753–764

Turgay K, Krause M, Marahiel MA (1992) Four homologous domains in the primary structure of GrsB are related to domains in a superfamily of adenylate-forming enzymes. Molecular Microbiology 6: 529–546

van Apeldoorn ME, van Egmond HP, Speijers GJA, Bakker GJI (2007) Toxins of cyanobacteria. Molecular Nutrition & Food Research 51: 7–60

Walden H (2010) Selenium incorporation using recombinant techniques. Acta Crystallographica Section D-Structural Biology 66: 352–357

Walsh CT, Chen H, Keating TA, Hubbard BK, Losey HC, Luo L, Marshall CG, Miller DA, Patel HM (2001) Tailoring enzymes that modify nonribosomal peptides during and after chain elongation on NRPS assembly lines. Current Opinion in Chemical Biology 5: 525–534

Wang JL, Li DD, Chen L, Cao W, Kong LL, Zhang W, Croll T, Deng ZX, Liang JD, Wang ZJ (2022) Catalytic trajectory of a dimeric nonribosomal peptide synthetase subunit with an inserted epimerase domain. Nat Commun 13

Weber T, Marahiel MA (2001) Exploring the Domain Structure of Modular Nonribosomal Peptide Synthetases. Structure 9: R3–R9

Winn M, Fyans JK, Zhuo Y, Micklefield J (2016) Recent advances in engineering nonribosomal peptide assembly lines. Nat Prod Rep 33: 317–347

Winn MD, Ballard CC, Cowtan KD, Dodson EJ, Emsley P, Evans PR, Keegan RM, Krissinel EB, Leslie AGW, McCoy A et al (2011) Overview of the CCP4 suite and current developments. Acta Crystallographica Section D 67: 235–242

Wu R, Reger AS, Lu X, Gulick AM, Dunaway-Mariano D (2009) The Mechanism of Domain Alternation in the Acyl-Adenylate Forming Ligase Superfamily Member 4-Chlorobenzoate: Coenzyme A Ligase. Biochemistry 48: 4115–4125

Yonus H, Neumann P, Zimmermann S, May JJ, Marahiel MA, Stubbs MT (2008) Crystal Structure of DltA IMPLICATIONS FOR THE REACTION MECHANISM OF NON-RIBOSOMAL PEPTIDE SYNTHETASE ADENYLATION DOMAINS. J Biol Chem 283: 32484–32491

Zeng XL, Huang M, Sun QX, Peng YJ, Xu XM, Tang YB, Zhang JY, Yang YL, Zhang CC (2023) A c-di-GMP binding effector controls cell size in a cyanobacterium. P Natl Acad Sci USA 120

